# *Zaxinone Synthase* overexpression modulates rice physiology and metabolism, improving growth and productivity under normal and low phosphate supply

**DOI:** 10.1101/2023.10.06.561198

**Authors:** Abdugaffor Ablazov, Muhammad Jamil, Imran Haider, Jian You Wang, Vanessa Melino, Moez Maghrebi, Gianpiero Vigani, Kit Xi Liew, Pei-Yu Lin, Guan-Ting Chen, Hendrik NJ Kuijer, Lamis Berqdar, Teresa Mazzarella, Valentina Fiorilli, Luisa Lanfranco, Xiongjie Zheng, Nai-Chiang Dai, Ming-Hsin Lai, Yue-Ie Caroline Hsing, Mark Tester, Ikram Blilou, Salim Al-Babili

## Abstract

The rice *Zaxinone Synthase (ZAS)* gene encodes a carotenoid cleavage dioxygenase (CCD) that forms the apocarotenoid growth regulator zaxinone. Here, we generated and characterized constitutive *ZAS*-overexpressing rice lines, to better understand *ZAS* role in determining zaxinone content and regulating growth and architecture. *ZAS* overexpression enhanced endogenous zaxinone level, promoted root growth and meristem size, and increased the number of productive tillers, leading to an up to 30% higher grain yield per plant. Hormone analysis revealed a decrease in strigolactone (SL) content, which we confirmed by rescuing the high-tillering phenotype through application of a SL analog. Metabolomics analysis revealed that *ZAS* overexpressing plants accumulate higher amounts of monosaccharide sugars, in line with transcriptome analysis. Moreover, transgenic plants showed higher carbon (C) assimilation rate and elevated root phosphate, nitrate and sulfate level, enhancing the tolerance towards low phosphate (Pi) and indicating a generally better nutrient uptake. Our study shows that *ZAS* regulates hormone homeostasis and a combination of physiological processes to promote growth and grain yield, which makes this gene an excellent candidate for sustainable crop improvement.

**Teaser:** *Zaxinone Synthase* overexpression modulates rice metabolism and physiology and improves growth and phosphate uptake.

## Introduction

Carotenoids are a class of tetraterpenes (C_40_) pigments, which are characterized by their long hydrocarbon chains containing a conjugated double-bond system (*1*). They are responsible for the vibrant colors seen in fruits and flowers and have an essential role in photosynthesis (*2-5*). Apocarotenoids, formed through the oxidative cleavage of double bonds in carotenoids, play a crucial role within the plant kingdom, as they serve as precursors for hormones and as pigments, aroma, scent constituents, and regulatory molecules (*6, 7*). Notably, abscisic acid (ABA) and strigolactones (SLs) are well-studied apocarotenoid hormones that play crucial roles in various aspects of plant growth, development, and adaptation (*8, 9*). The formation of apocarotenoids is primarily catalyzed by a family of carotenoid cleavage dioxygenases (CCDs), which exhibits divergent cleavage properties leading to the formation of various apocarotenoids with unique features and functions (*10, 11*). The genome of *Arabidopsis thaliana* encodes nine CCD members, which are designed as CCD1, CCD4, CCD7, CCD8, and five 9-*cis*-epoxycarotenoid cleavage dioxygenases (NCED2, 3, 5, 6, and 9). CCD1 and CCD4 enzymes possess a broader substrate specificity, allowing them to cleave a diverse range of carotenoids and apocarotenoids, which results in various volatiles and/or apocarotenoid pigments (*7*). CCD7 is a SL biosynthetic enzyme cleaving 9-*cis*-β-carotene at positions 9, 10, or 9′,10′ to produce 9-*cis*-β-apo-10-carotenal and β-ionone (*12*). Then, CCD8 mediates this intermediate into carlactone (CL), the precursor of SLs, by cleaving at positions 13, 14 of 9-*cis*-β-apo-10-carotenal (*12*). NCEDs are involved in ABA biosynthesis, which cleaves the 11, 12 (11′, 12′) double bond of 9-*cis*-epoxy carotenoids to yield xanthoxin, the precursor of ABA (*13, 14*). Recent studies revealed a bypass in ABA biosynthesis, leading to non-epoxydated β-apo-11-carotenoids to xanthoxin (*15*).

ZAS, a recently discovered member of the CCD family, cleaves apo-10′-zeaxanthinal, at the C13–C14 double bond, leading to zaxinone (apo-13-zeaxanthinal) (*16*). Zaxinone is a growth regulator and an apocarotenoid hormone candidate, which is required for normal rice growth and development and negatively regulates SL biosynthesis (*16, 17*). Corresponding *ZAS* loss-of-function mutant in rice exhibited reduced zaxinone content in the roots, accompanied by severe growth defects, such as reduced root and shoot biomass, lower tiller number, and increased SL level (*16*). Exogenous application of zaxinone partially rescued the growth defects in *zas* mutant and also promoted root growth in wild-type seedlings, and repressed SL biosynthesis (*16*). Moreover, exogenous application of zaxinone mimics led to enhanced growth and productivity of major agricultural crops, such as capsicum, potato, strawberry, and other crops in the field conditions (*18, 19*).

Interestingly, there are no *ZAS* orthologues in non-mycorrhizal species, such as Arabidopsis, which indicates a role of this gene in the arbuscular mycorrhizal (AM) symbiosis (*16, 20*). This association between AM fungi and host plants plays a crucial role in providing essential minerals such as phosphorus (P) and nitrogen (N) to the plant, while the AM fungi receive carbohydrates and lipids in return (*21*). In fact, the *zas* mutant exhibited a lower level of AM colonization compared to wild-type plants. Conversely, overexpression of *ZAS* under the control of the *OsPT11* promoter, which is known to be active in arbusculated cells, resulted in increased mycorrhization (*22*). Furthermore, *ZAS2* (homologue of *ZAS*) knock-out plants showed impaired AM colonization (*23*). These findings indicate that *ZAS* gene family members play an essential role in ensuring the successful establishment of AM symbiosis in rice.

Previous studies demonstrate that rice requires a functional *ZAS* gene and a certain level of zaxinone in its roots to maintain normal growth and symbiosis with AM fungi. However, the effect of increasing zaxinone biosynthesis on rice growth, physiology, and architecture remained elusive. In this study, we addressed this question by generating and characterizing constitutive *ZAS*-overexpressing rice lines.

## Results and Discussion

### Overexpression of *ZAS* enhances zaxinone level, increases number of productive tillers and total grain weight per plants

We set out to explore the effect of increasing *ZAS* expression on rice growth and architecture, particularly tillers number and root size, and on its productivity. To achieve this, we generated transgenic rice plants overexpressing *ZAS* under the control of the constitutive *CaMV 35S* promoter (fig. S1A). For further studies, we selected three independent transgenic lines (designed as *OX1, OX6*, and *OX9*) with an around 90- and 1000-fold higher expression level, compared to wild type, in shoots and roots, respectively (Fig. 1A). Selected lines were grown until the T4 generation, with transgenicity confirmed through growth on hygromycin-containing media and reverse transcription-quantitative polymerase chain reaction (RT-qPCR) analysis. Southern blot analysis using the *hygromycin phosphotransferase* (*HPT*) probe confirmed the presence of a single copy of the inserted *T-DNA* (fig. S1B). Zaxinone quantification of hydroponically grown *ZAS*-overexpression plants showed a significant increase in the range of 20 and 85% in shoot and root tissues, respectively (Fig. 1B), suggesting the role of ZAS in determining internal zaxinone levels. Next, we characterized the lines during the maturing stage under greenhouse (GH) conditions. Compared to wild type, the overexpression lines developed significantly more tillers (∼27%), which were mostly productive at the heading stage (Fig. 1, C and E). With respect to panicles and grains, we observed a slight reduction in panicle length and number of grains per panicle, compared to wild type, with the exception of *OX6* (Fig. 1, I, K and L). We also detected a slight decrease in grain size (Fig. 1, J, M and N), but no significant change in the 1000-grains weight (Fig. 1H). In summary, *ZAS*-overexpressing lines produced about 30% higher grain weight per plant (Fig. 1, D and F). The total above-ground biomass also increased by an average of 20-25% (Fig. 1G), with a tendency to reduced height (fig. S2A). With respect to the flowering time, we did not observe a significant difference to wild type plants (fig. S2B). The possible reason for the decrease in grain and panicle size could be attributed to the enhanced shoot growth and increased tillers in *ZAS* overexpression lines. This excessive growth might lead to a dilution of carbohydrates or other essential resources during the plant’s vegetative stage. Consequently, the transgenic lines might have limited resources compared to the wild-type plants when it comes to allocating for the development of grains and panicles. However, overexpression of *ZAS* significantly enhanced the number of productive tillers, more than compensating for the reduction in the number and size of grains, ultimately resulting in higher yield under GH conditions.

**Fig. 1.**
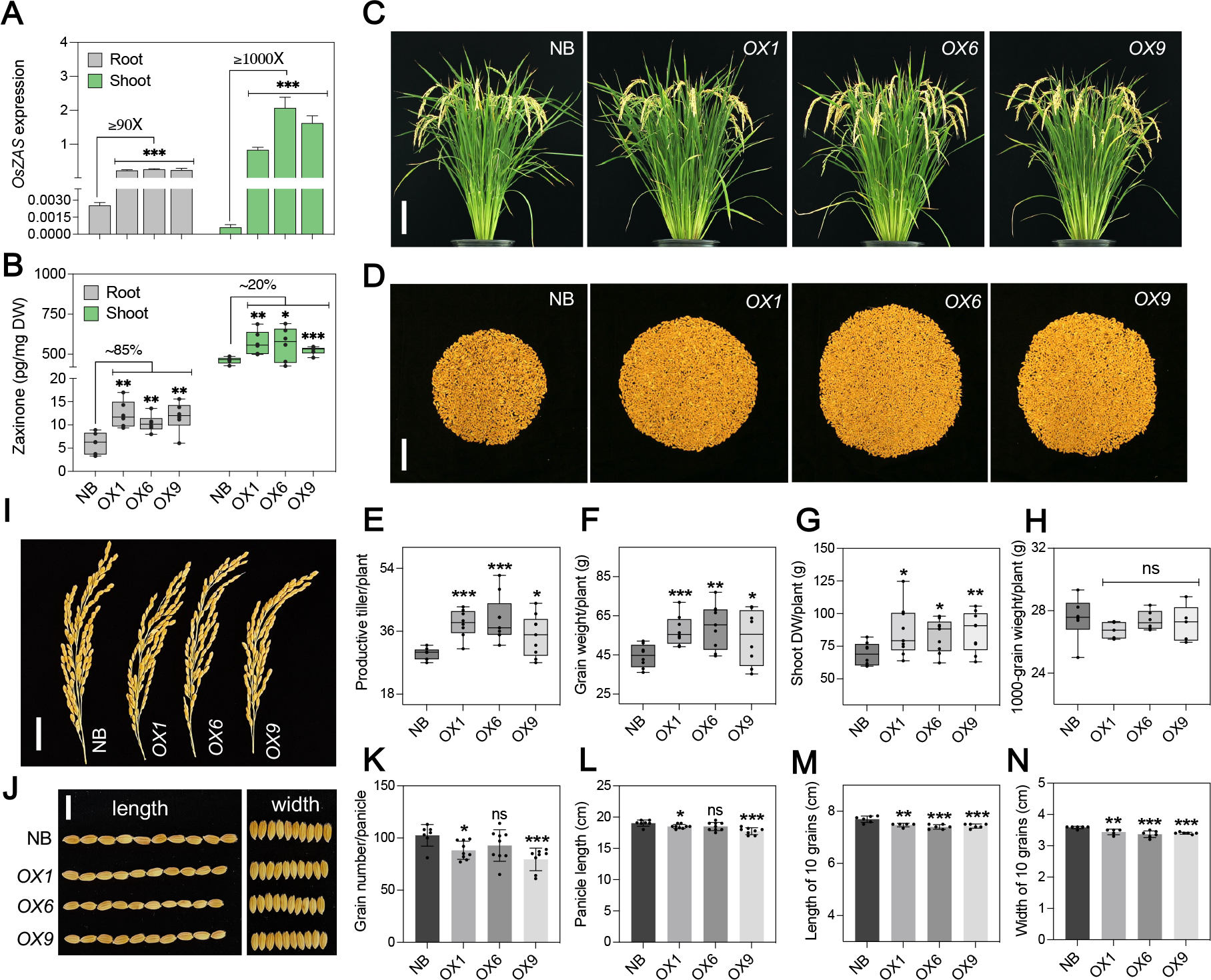
Overexpression of *ZAS* promotes rice tillering and yield. (**A**) Normalized expression value of *ZAS* analyzed with qRT-PCR in transgenic rice lines roots and shoots, respectively. Data represent mean ± SD (*n* ≥*3*). (**B**) Quantification of zaxinone content in *ZAS* overexpression lines in the shoots and roots, respectively. Data represent mean ± SD (*n* ≥*4*). (**C**) Plant architecture of Nipponbare wild type (WT) and transgenic lines overexpressing *ZAS* (*OX1, OX6, OX9*) at maturing stage. Bar graph is 10 cm. (**D**) Presentation of total grains per plant. Bar graph is 5 cm. (**E**) Productive tiller number per plant. (**F**) Grain weight per plant. (**G**) Shoot dry weight (DW) per plant. (**H**) 1000-grain weight per pant. (**I**) Representation of main panicles per plants. Bar graph is 5 cm. (**J**) Length and width of 10 seeds from selected replicas. Bar graph is 1 cm. (**K**) Grain number per main panicle. (**L**) Length of main panicle. (**M**) Length of 10 grains separated from main panicle. (**N**) Width of 10 grains separated from main panicle. Boxes in boxplots represent the median, first and third quartile. The minimum and maximum values are shown with the length of the whiskers. Dots in the boxplots represent the biological replicates. Values in (K–N) are means ± SD (*n* ≥ 9). Student’s *t* test used for the statistical analysis (\**P* < 0.05;

### ZAS promotes the tillering by suppressing the strigolactone (SL) biosynthesis

Tillering is a treat controlled by different growth conditions and regulatory genes and is determined by phytohormones (*24-27*). Strigolactones (SLs) are a major inhibitor of axillary bud outgrowth (*25, 28, 29*). Mutants disrupted in SL biosynthesis or perception, such as *ccd7* (*d17*), *ccd8* (*d10*) or *d14*, are characterized by a high-tillering and dwarf phenotype, accompanied by a significant reduction in yield due to the dominance of non-productive tillers (*9*). Previously, we showed that zaxinone is a negative regulator of SL biosynthesis in rice. Loss-of-function of *ZAS* and its orthologue *ZAS2* led to increased SL content and a reduction in the number of tillers (*16, 23*). Based on the phenotypes observed in *zas* mutants, we hypothesized that high tillering in *ZAS-*overexpression lines might be due to SL deficiency. To check this assumption, we quantified SLs by using LC/MS; however, their content under optimal growth conditions, *i*.*e*. sufficient Pi supply, was below detection limit. Therefore, we used the more sensitive *Striga* seed germination bioassay. Results obtained showed a drastic decrease in *Striga* seed germinating activity of root extracts of *ZAS*-overexpression lines, compared to wild-type (Fig. 2A), indicating a lower SL content. We obtained similar results with root exudates that showed around 50% reduction in *Striga* seed germinating activity (Fig. 2B). Consistent with the lower SL content, the transcript level of two main SL biosynthetic genes. i.e. *CCD7* and *CCD8*, was lower in *ZAS-*overexpression lines, compared to wild-type (Fig. 2, C and D). To further confirm that the high-tillering phenotype was a result of a decrease in SL content, we applied the SL analog methyl phenlactonoates 3 (MP3) (*30*) to soil-grown *ZAS*-overexpression lines. Treatment with MP3 inhibited the growth of the second tiller in the *ZAS*-overexpression lines, but not in wild type (Fig. 2, E and F). The inhibition observed upon MP3 treatment indicates that SL deficiency in *ZAS*-overexpression lines is a reason for their increased tiller number. Taken together, our findings suggest a correlation between a higher content of zaxinone in *ZAS*-overexpression lines and a reduction in SL biosynthesis and consequences thereof (Fig. 2G).

**Fig. 2.**
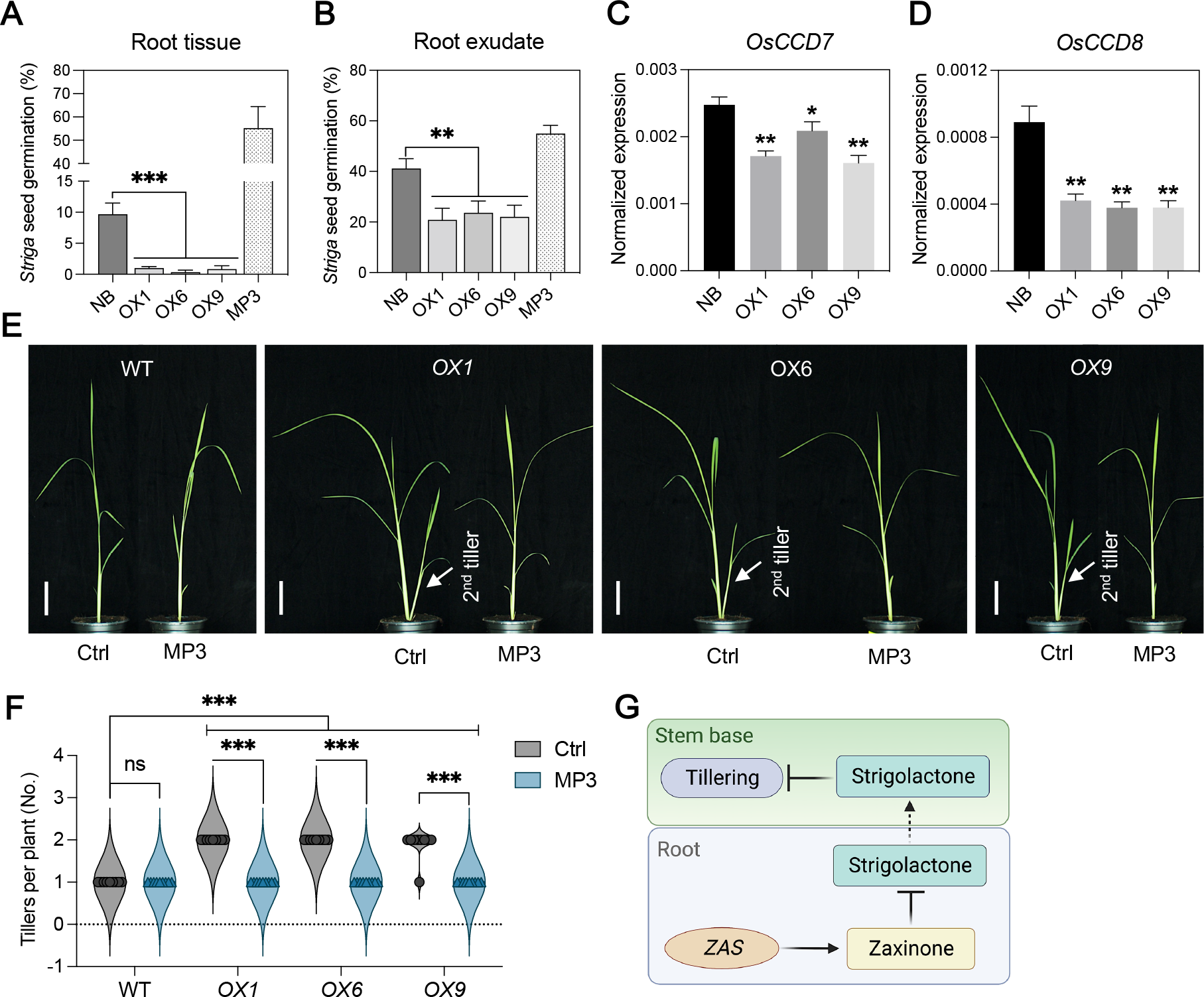
ZAS regulates tiller development by repressing the SL biosynthesis in rice. (**A**) and (**B**), SL quantification in roots tissue and exudates of wild-type and *ZAS* overexpression lines with the *Striga* germination assay. 1 μM of MP3 (SL mimic) used as a control. Data represent mean ± SD (*n* ≥*5*). (**C**) and (**D**), Two main SL biosynthetic genes (*CCD7* and *CCD8*) expression determined by qRT-PCR in the root tissue of wild-type and *ZAS* overexpression lines. Data represent mean ± SD (*n* ≥*3*). (**E**) Picture of tillering rescue experiment of *ZAS* overexpression transgenic lines which performed in the soil by applying MP3 (5 μM) for one week. Bar graph is 5 cm. (**F**) Quantification of tiller number in wild-type and ZAS overexpression lines after MP3 application. Values in (F) are means ± SD (*n* = 9). Student’s *t* test used for the statistical analysis (\**P* < 0.05; \*\**P* < 0.01; \*\*\**P* < 0.001). (**G**) Proposed model that explains how *ZAS* regulates tillering in rice.

### *ZAS* overexpression enhances the root growth by regulating the meristem activity

Besides being an anchorage in soil, roots are crucial for proper nutrient and water uptake, which ultimately determines the growth and overall development of plants. Overexpression of *ZAS* significantly increased the growth and number of crown roots in both solid (1/2 MS, 0.4% agarose) and liquid (hydroponic) medium, as determined at different time points in comparison to wild-type plants (figs. S3A and B). Moreover, the *ZAS*-overexpression lines showed a significantly enhanced root length and biomass (Fig. 3, A to D). To gain insights at cellular level, we examined the meristem size in root tips, which indicates the rate of cell division, using bright-filed microscopy. *ZAS*-overexpression lines displayed larger root meristem size, compared to the wild-type (Fig. 3, E and F). Confirming increased meristem size, the number of meristematic cells in *ZAS*-overexpression lines was higher than in wild-type (Fig. 3G). These data indicate that overexpression of *ZAS* promotes root elongation by enhancing the meristem activity and cell division rate. Our findings are consistent with results obtained by exogenous zaxinone treatment that led to an increase in meristem size and cell number (*31*) and suggest that the higher level of zaxinone in transgenic lines can stimulate the cell division rate, thereby promoting root growth and development.

**Fig. 3.**
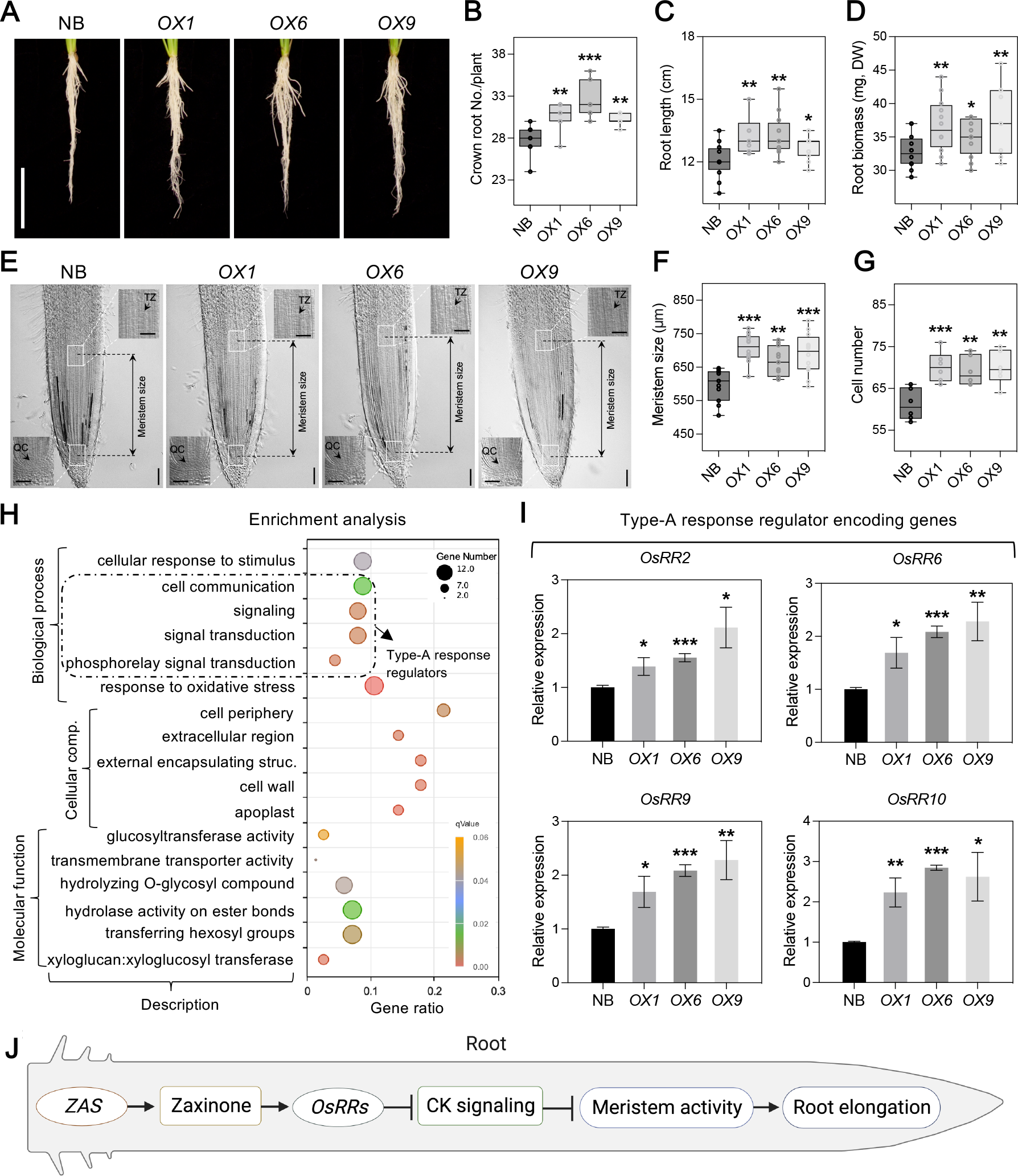
ZAS enhances root growth through regulating meristem activity. (**A**) Root phenotype of *ZAS* overexpression lines grown in the hydroponics medium for three weeks. (**B**) Crown root numbers (per plant) of *ZAS-*overexpression lines. (**C**) Root length of *ZAS* overexpression lines. (**D**) Root biomass (DW, dry weight) of *ZAS-*overexpression lines. (**E**) Root tips of 5-day-old *OsZAS* transgenic and wild-type seedlings were examined under light microscope with 10X and 20X. The distance between the quiescent center (QC) and transition zone (TZ) represents the size of meristem (between two dash lines). Horizontal and vertical bars in the pictures represent 50 and 100 μm, respectively. (**F**) The meristem size was measured in the root tips of *ZAS-*overexpression and wild-type with 10X (n=11-12). (**G**) The number of central cylinder cells was counted in the meristem region of *ZAS-* overexpression and wild-type root tips with 20X (n=6). (**H**) Gene Ontology (GO) term analysis of up-regulated genes in roots of *ZAS* overexpression (*OX6*) transgenic plants. GO terms with corrected *P*-value ≤ 0.05 were considered significantly enriched by up-regulated genes. (**I**) Relative expression level of rice type-A cytokinin response regulator encoding genes in three-week-old rice seedlings (*n* ≥*3*). (**J**) Proposed model about how overexpression of *ZAS* improves the rice root growth. Boxes in boxplots represent the median, first and third quartile. The minimum and maximum values are shown with the length of the whiskers. Dots in the boxplots represent the biological replicates. Student’s *t* test used for the statistical analysis (\**P* < 0.05; \*\**P* < 0.01; \*\*\**P* < 0.001; ns, not significant).

The regulation of root meristem activity and cell proliferation is predominantly controlled by the plant hormones auxin and cytokinins (CKs) (*32-35*), which have opposing roles in root development, with auxin being a promoter of cell proliferation while CKs negatively impacts meristem size. Therefore, we hypothesized that *ZAS* overexpression may enhance cell division rate by modulating auxin/cytokinin homeostasis. To test this hypothesis, we measured the levels of the auxin indole-3-acetic acid and active CKs (Isopentenyl adenine, *trans*- and *cis*-Zeatin) in the roots of *ZAS*-overexpression lines. Compared to wild type, we did not detect a change in auxin or *cis*-zeatin (cZ) content (figs. S4, A and B), but observed a moderate decrease in the level of isopentenyl adenine (iP) and a significant enhancement in *trans*-zeatin (tZ) content (fig. S4B). We also determined the level of other hormones. We did not observe a change in abscisic acid (ABA) and gibberellic acid (GA) level, but detected an increase in salicylic acid (SA) content in *ZAS-OX* lines, compared to wild-type (figs. S4, C to E). In conclusion, quantification of hormones did not provide a clear explanation for the root phenotype. Therefore, we assumed that the overexpression of *ZAS* might influence the signaling pathways of these hormones. To test this hypothesis, we carried out RNAseq experiment on roots of the *ZAS*-overexpression line *OX6*. There was no significant enrichment in GO (Gene Ontology) terms related to auxin signaling, but we observed an enrichment of CK signaling associated pathways, including the phosphorelay signal transduction and cell communication (Fig. 3H). In particular, there was an up-regulation of genes encoding type A response regulators (RRs) that act as inhibitor of CK signal transduction in plants (*36-38*). As confirmed by qRT-PCR, transcript level of *OsRRs* (*OsRR2, 6, 9, 10*) was significantly higher in *ZAS*-overexpression lines (Fig. 3I), which may explain the observed root phenotype. Indeed, previous studies demonstrated that overexpression of *OsRR2* increased the number of crown roots (*39*), while *OsRR3-* and *OsRR5*-overexpressing lines exhibited reduced sensitivity to exogenous CK application (*40*), indicating their role in regulating CK signaling and root development. To confirm the reduced CK-sensitivity in the roots of our transgenic lines, we germinated rice *OX1* and wild-type seeds on 1/2 MS (Murashige and Skoog) medium in the presence or absence of the CK analog 6-Benzylaminopurine (BA, 1 μM concentration). Supporting our hypothesis, *OX1, ZAS*-overexpressing plants showed, upon BA treatment, around 45% lower root growth inhibition, compared to wild-type (figs. S5, A and B). Taken together, our findings indicate that *ZAS* likely enhances root elongation by increasing *OsRRs* activity and thus inhibiting CK signaling, which results in a higher cell division rate in root meristem (Fig. 3J).

### Overexpression of *ZAS* leads to an increase of monosaccharide sugars accumulation and enhances the rate of carbon (C) assimilation

Sugars play a significant role as universal carbon source for building cellular components in living organisms, in addition to their function in providing the energy required for growth and development. We previously demonstrated that exogenous zaxinone application promoted rice growth by enhancing sugar metabolism (*31*). Therefore, we hypothesized that increased zaxinone content that is caused by overexpressing *ZAS* may improve rice growth through enhancing sugar accumulation. To test this hypothesis, we measured the levels of different sugars in both shoot and root tissues of the *ZAS-* overexpressing lines *OX1* and *OX6*, using gas chromatography–mass spectrometry (GC– MS). Our results showed a significant increase in different monosaccharides, such as glucose, fructose, and myo-Inositol, in roots and shoots of the overexpression lines (Fig. 4A). However, disaccharide sugars, such as sucrose, maltose, galactinol, and trehalose, remained generally unchanged or decreased in overexpression lines (fig. S6), indicating a dynamic sugar metabolism where disaccharide sugars might be quickly hydrolyzed into simple sugars or utilized for building up polysaccharides. Analysis of hexose phosphates, such as Glucose-6-phosphate (G6P) and Fructose-6-phosphate (F6P), revealed a significant increase in both roots and shoots, with exception of Glucose-1-phosphate (G1P) that was slightly reduced in the shoots of the two *ZAS*-overexpressing lines (Fig. 4B). Previous studies suggest that sugar phosphates act as signaling molecules repressing the activity of SnRK1 (SUCROSE-NON-FERMENTING1-RELATED KINASE1), a key negative regulator of plant growth that promotes catabolic processes while inhibiting energy-using anabolic processes (*27, 41-43*). It could be speculated that *ZAS* promotes rice growth through suppressing SnRK1 activity by increasing the levels of sugar phosphates. Consistent with the sugar measurements, Gene ontology (GO) enrichment analysis of up-regulated genes, deduced from our RNA sequencing (RNA-Seq) experiment, revealed a significant increase in terms related to sugar metabolisms (carbohydrates, polysaccharides, disaccharides, and trehalose) in *ZAS* overexpression lines (Fig. 4C). Taken together, our findings suggest that *ZAS* promotes sugar accumulation, particularly of monosaccharides, likely through increasing zaxinone content.

**Fig. 4.**
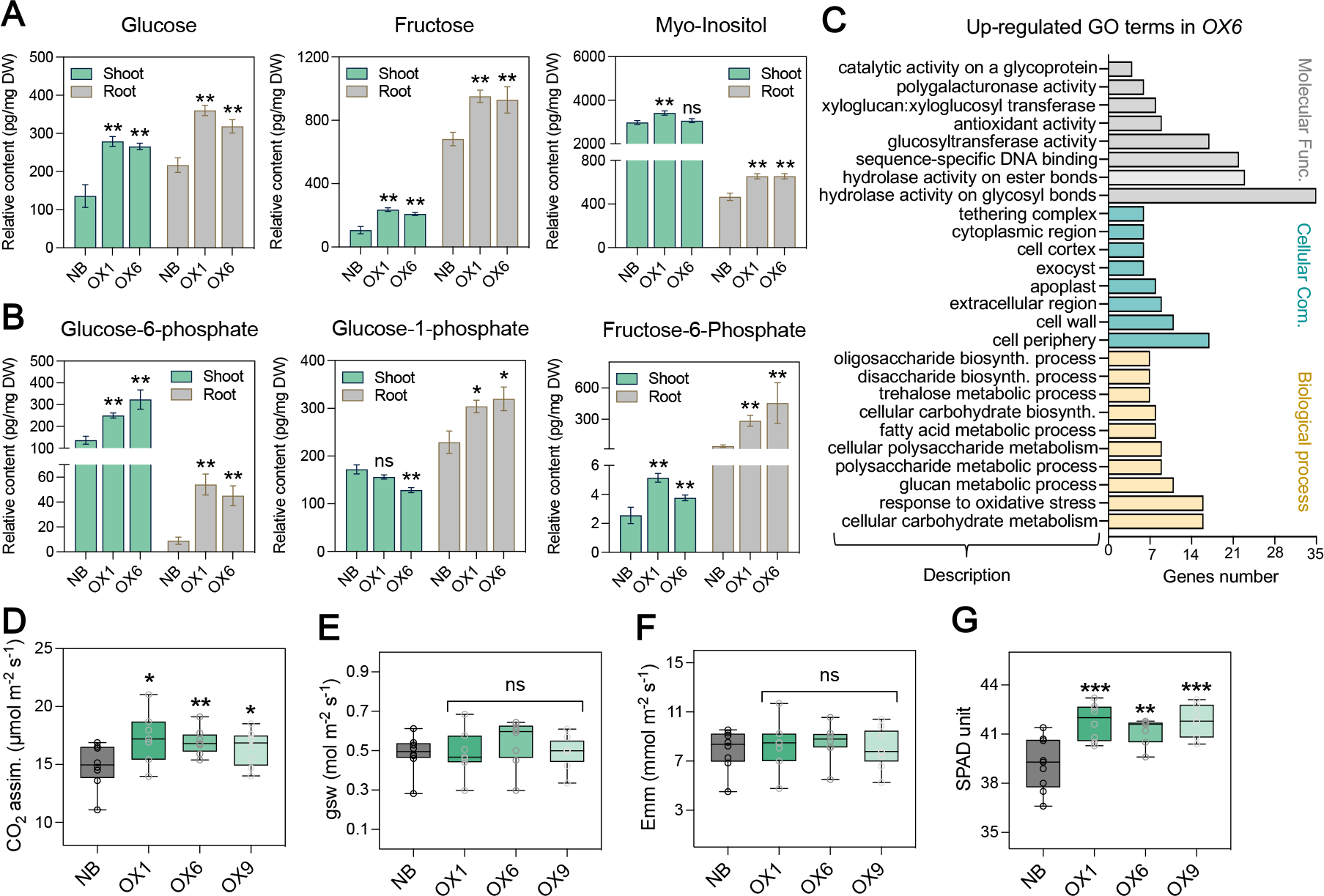
ZAS enhances monosaccharide sugars accumulation and carbon dioxide (CO_2_) assimilation. (**A**) SL Monosaccharaide sugars were quantified in both root and shoot of three-week-old rice seedlings grown in hydroponics. (**B**) Hexose sugars were quantified in both root and shoot of three-week-old rice seedlings. Values (A-B) represent mean ± SD (*n* ≥*5*). (**C**) Gene Ontology (GO) term analysis of up-regulated genes in shoots of *ZAS* overexpression (*OX6*) from RNA-seq. GO terms with corrected *P*-value ≤ 0.05 were considered significantly enriched by up-regulated genes. (**D**) Carbon dioxide (CO_2_) assimilation rate per unit of projected leaf area. (**E**) Stomatal conductance to water vapor (gsw) per unit of projected leaf area. (**F**) Transpiration rate (Emm) per unit of projected leaf area. (**G**) The relative chlorophyll content was measured using SPAD-502Pus (Konica Minolta). Values (D-G) represent mean ± SD (*n* ≥*8*). Boxes in boxplots represent the median, first and third quartile. The minimum and maximum values are shown with the length of the whiskers. Dots in the boxplots represent the biological replicates. Student’s *t* test used for the statistical analysis (\**P* < 0.05; \*\**P* < 0.01; \*\*\**P* < 0.001; ns, not significant).

Based on the increased monosaccharides level, we assumed that overexpression of the *ZAS* gene may enhance the photosynthetic activity. To check this assumption, we analyzed C assimilation and transpiration rate, stomatal conductance and relative chlorophyll content of the youngest fully expanded leaves of four-week-old rice plants, using the LICOR-6800 Gas Exchange system and SPAD-502 Chlorophyll meter. Results obtained indicated that *ZAS*-overexpression lines have a higher net rate of CO_2_uptake and relative chlorophyll content per unit of projected leaf area, compared to the wild-type (Fig. 4, D and G). However, we did not observe significant difference in the stomatal conductance or transpiration rate between the *ZAS*-overexpression plants and wild-type (Fig. 4, E and F). From these findings, we conclude that *ZAS* fosters photosynthesis by facilitating C assimilation and increasing chlorophyll content, which can explain the higher sugar accumulation in overexpression lines.

### *ZAS* overexpression improved major macro-nutrients uptake in roots

Considering the overall improved performance of *ZAS-OX* lines and increased root growth, we assumed that *ZAS* overexpression might improve nutrient uptakes. To test this assumption, we quantified phosphates (PO_4_^3−^), nitrate (NO_3_^−^), and sulfate (SO_4_^2−^) in roots and shoots of *ZAS* overexpressing lines. We observed a striking increase in phosphate (∼60%) and sulfate (∼90%) content in roots of the overexpression lines, compared to wild type (Fig. 5A; fig. S7A). We also detected an increase in nitrate content, but to a less pronounced extent (fig. S7B). However, we did not observe any significant change in all measured nutrients in shoots of *ZAS-*overexpression lines (Fig. 5A; figs. S7, A and B). It is possible that the reason for the higher levels of nutrients found in the roots, rather than the shoots, of the *ZAS*-overexpression lines could be a decrease in nutrient utilization within the roots. This phenomenon may be a result of altered physiological process caused by the overexpression of the *ZAS* gene. Further investigation is needed to fully understand the mechanisms behind this observation. In summary, our data suggest that the overexpression of *ZAS* enhanced the uptake of major nutrients, particularly sulfate and phosphate. The increase in phosphate content may explain why overexpression of *ZAS* lead to a decrease in SL level, given the inverse relationship between phosphate content and SL biosynthetic rate in many plant species, including rice (*24, 25, 27, 44-46*).

**Fig. 5.**
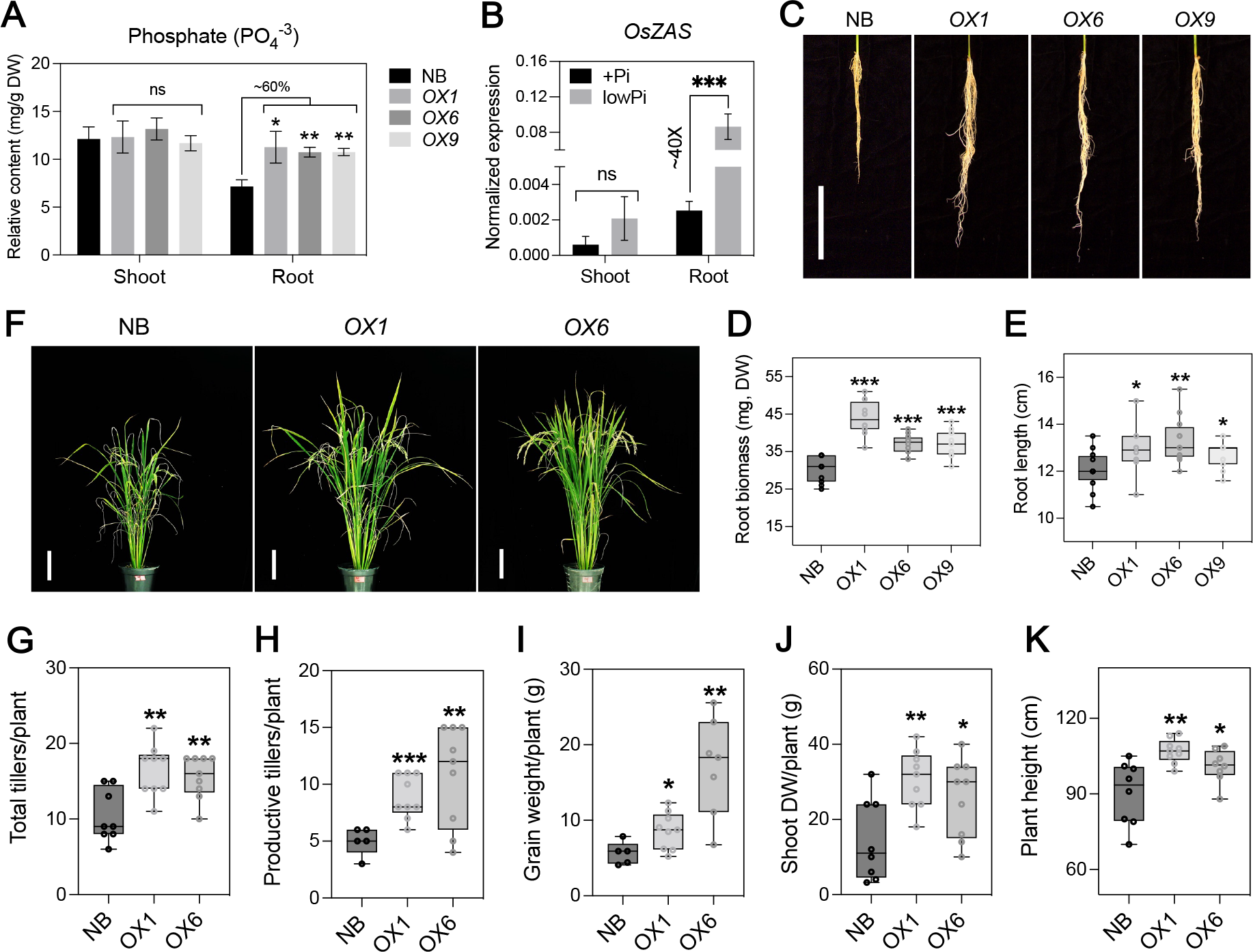
ZAS enhances phosphate (Pi) uptake and low Pi tolerance. (**A**) Phosphate content was quantified in both roots and shoots of wild type (NB) and *ZAS* overexpression lines grown for three weeks in hydroponics (mean ± SD; *n* ≥*4*). (**B**) Normalized expression level of *OsZAS* in three-week-old rice seedlings grown under +Pi (400 μM K_2_HPO_4_.2H_2_O) and low Pi (4 μM K_2_HPO_4_.2H_2_O) conditions for two-weeks (mean ± SD; *n* ≥*3*). (**C**) Root phenotype of *ZAS-*overexpression lines grown under low Pi conditions for two-weeks after one-week pretreatment growth. (**D**) Quantification of root biomass (DW, Dry Weight) shown in (C). (**E**) Measurement of root length shown in (C). Values represented in d and e are means ± SD (*n* ≥ 11). (**F**) Plant architecture of wild type and transgenic lines overexpressing *ZAS* at maturing stage grown in low Pi soil under GH conditions. (**G-K**), Total tiller number per plants (**G**), productive tiller number per plants (**H**), grain weight per plants (**I**), shoot biomass (**J**), plant height (**K**) of NB and *ZAS*-overexpression lines (*OX1, OX6*) represented in (**F**). Values in (G-K) are means ± SD (*n* ≥ 5). Boxes in boxplots represent the median, first and third quartile. The minimum and maximum values are shown with the length of the whiskers. Dots in the boxplots represent the biological replicates. Student’s *t* test used for the statistical analysis (\**P* < 0.05; \*\**P* < 0.01; \*\*\**P* < 0.001; ns, not significant).

### *ZAS* overexpression enhances the growth and productivity under low Pi tolerance

Supporting the role of ZAS in improving Pi uptake, a recent study revealed the presence of the Pi starvation-responsive binding site (P1BS) in the *ZAS* promoter, and that the expression of this gene is transactivated by PHOSPHATE STARVATION RESPONSE 2 (PHR2), a key regulator of Pi signaling and arbuscular mycorrhizal (AM) symbiosis, upon Pi deficiency in rice (*47*). Consistently, the expression of *ZAS* significantly increased (by about 40-fold) under low Pi conditions in roots, but remained unaffected in shoots (Fig. 5B). These data indicate that *ZAS* could enhance low Pi tolerance in rice. To test this hypothesis, we evaluated the performance of our overexpression lines under low Pi conditions, when grown under controlled conditions in both hydroponic medium and soil. The overexpression of *ZAS* significantly improved root growth, as evidenced by increased root biomass and elongation, compared to the wild type (Fig. 5, C to E). Moreover, overexpression lines grown in low Pi soil under GH conditions maintained their high-tillering phenotype observed under Pi-sufficient conditions (Fig. 5, F to H). Furthermore, grain weight and above ground biomass were significantly higher in *ZAS*-overexpression lines, compared to wild-type (Fig. 5, I and J). Interestingly, *ZAS*-overexpression lines were taller than wild-type under low Pi conditions, in contrast to what we observed under normal conditions (Fig. 5, F and K). Overall, these findings suggest that ZAS enhances Pi uptake and hence improves the tolerance to low Pi-conditions.

### Overexpression of *ZAS* did not affect AM fungi colonization

Recently, we showed that overexpression of *ZAS* under the control of the *OsPT11* promoter, that is known to be active in arbusculated cells led to increased mycorrhization (*22*). Therefore, we investigated the mycorrhization capability of our constitutive *ZAS* overexpressing lines. As shown in (figs. S8, A and B), we did not detect any significant alteration in AM colonization. This inconsistence could be explained by the different activity of the promoters used to generate the overexpressing lines. Indeed, *CaMV 35S* is a strong constitutive promoter which exhibits high transcriptional activity and stability compared to *OsPT11* promoter, that is exclusively active in arbuscule-containing cells and in rice root apexes. This is mirrored in zaxinone content since in the root of *35S prom::OsZAS* lines, zaxinone reaches a significant increment (∼85%), while in the root of *OsPT11prom::OsZAS* lines zaxinone increment was lower (∼45%) (*22*). Moreover, contrarily to *35S prom::OsZAS* lines, *OsPT11prom::OsZAS* lines displayed an increment of SLs content compared to WT, which is in agreement with the increased mycorrhization level observed in these lines.

## Conclusion

In summary, we show here that constitutive overexpression of *ZAS* caused a novel combination of effects, which promotes rice growth and increases its productivity by enhancing the number of productive tillers and improving root growth, nutrient uptake and photosynthesis, without having negative impact on mycorrhization. Importantly, *ZAS* overexpression decreased the level of SLs, which explains the increased the high-tillering and may be a result of better Pi uptake. The fact that *ZAS* overexpressing lines, in contrast to SL-deficient mutants, developed productive tillers may be a result of an increased shoot biomass, photosynthetic capacity and nutrient uptake, which make the resources available, which are required for the observed significant increase in grain productivity. The increase in sugar content also supports this assumption. Although we are still at the beginning of understanding zaxinone biology, our study demonstrates the importance of *ZAS* as a target for sustainable enhancement of rice performance and for reducing the demand for Pi fertilizers. In addition, our findings mostly align with the exogenous zaxinone effect on rice growth and development. Thus, our study offers an alternative strategy to enhance the rice growth performance and productivity by manipulation of *ZAS* expression, negating the need for exogenous zaxinone application.

## Materials and Methods

### Plant material and growth conditions

In this study, we utilized *Oryza sativa* L. cv. Nipponbare rice as experimental model. Rice seeds were first incubated in 2% sodium hypochlorite (v/v) for 15 min and rinsed five times in sterile double-distilled water. Seeds were then germinated on ½ MS (0.4% Agarose) medium for two days under dark conditions at 30°C. After two days, seeds were moved to a growth chamber (Conviron) with 28°C Day/22°C night rhythm and a photoperiod of 12-hour light/12-hour dark. The photon irradiance was set at 500 μmol. m−2.s^−1^ and the relative humidity at 70%. Then, the 7-day-old seedlings were hydroponically grown in Hoagland’s nutrient solution for two weeks, according to (*23*). For metabolite and molecular analysis, root and shoot samples were harvested and stored at -80°C until analysis. For the low Pi hydroponics experiment, 7-day-old seedlings were subjected low Pi (4 μM K_2_HPO_4_x2H_2_O) deficiency for two weeks, and root and shoot phenotypes were recorded. In order to characterize the yield-related phenotypes of *ZAS* overexpression transgenic rice lines, 7-day-old seedlings were transferred into soil-filled pots in a Greenhouse (GH) with specific conditions (temperature 28°C Day/25°C night; photoperiod 12-h-light/12-h-dark; light intensity ∼800 μM m^−2^s^−1^ and 70% relative humidity). Rice plants were watered twice a week with modified half-strength Hoagland’s nutrient solution. For the low Pi soil experiment, a single plant was grown in a 2-liter pot filled with 1 kg of stender soil (153 mg Pi/kg as P_2_O_5_) and 1-liter of sand mixture. The amount of Pi was defined and calculated as low Pi based on (*48*). The rice plants were supplemented twice a week with -Pi (without K_2_HPO_4_x2H_2_O) modified half-strength Hoagland’s nutrient solution. At maturity, agronomic traits were recorded. The AM fungi experiment was performed as described by (*49*).

### Striga hermonthica seed germination bioassay and SL quantification

Striga seeds were preconditioned following the procedure described by (*50*). Root extracts and exudates were prepared and applied as described by (*23*). Then, SL containing samples were applied on pre-conditioned Striga seeds and kept at 30°C for 24 h in the dark. Moreover, the SL analog MP3 (1 μM) and MillQ water were applied as positive and negative control, respectively. Germinated and non-germinated seeds were scanned with a light microscope and analyzed by using the software SeedQuant (*51*). SLs in rice root exudates were enriched as described by (*52*). SLs in root tissues were extracted following the protocol described by (*16*). SL samples were analyzed by using UHPLC-Triple-Stage Quadrupole Mass Spectrometer (TSQ-Altis) with parameters as mentioned in (*52*).

### Tillering rescue bioassays

7 days-old seedlings of the Nipponbare wild type and *ZAS* overexpressing lines (*OX1, OX6, OX9*), were planted in soil (500 mL plastic pots) supplemented with the synthetic SL analog MP3 (5 µM) and acetone (0.05%) as a control. MP3 solution was applied every three days for two weeks and tillers per plant were counted and pictured with digital camera.

### Quantitative analysis of zaxinone

Zaxinone content was quantified using the previously described protocol (*16*). Approximately 20 mg of lyophilized rice samples were extracted twice with 2 mL of ethyl acetate containing 2 ng of D3-zaxinone (customized synthesis; Buchem B.V., Apeldoorn, The Netherlands). After that, samples were sonicated for 15 minutes in an ultrasonic bath and centrifuged at 3800 rpm for 8 minutes at 4°C. The two supernatants were combined and then dried under vacuum. The dried extract was dissolved in 100 µL of ethyl acetate and 2 mL of hexane before being purified. Silica gel SPE column (500 mg/3 mL) were used for purification, pre-conditioned with 3 mL of ethyl acetate and 3 mL of hexane. After washing with 3 mL of hexane, zaxinone was extracted using 3 mL of ethyl acetate and evaporated to dryness under vacuum. The residue was then re-dissolved in a mixture of 120 μL acetonitrile:water (25:75, v:v) for zaxinone and filtered through a 0.22 μm filter for UHPLC-Triple-Stage Quadrupole Mass Spectrometer (TSQ-Altis) with parameters as described in (*52*).

### Meristem size measurement and cell number quantification

Seeds of Wild type (NB) and *ZAS*-overexpression lines were placed on a 0.8% Agarose (1/2MS) plate and allowed to germinate in complete darkness at 28°C for 2 days. The plates were later moved to a growth chamber (Conviron) and kept under specific conditions: a 12-hour light/12-hour dark cycle at a temperature of 28°C during the day and 22°C at night, with photon irradiance set at 500 μmol. m−2.s^−1^, and a relative humidity of 70%. After 96 hours, roots were cut, and seedlings were subjected to chloral hydrate solution for five days at a temperature of 4°C. Root tips were then fixed onto a slide and photographed under a light microscope with 10X and 20X magnification. Using the ImageJ software, Meristem size and cell number were analyzed, using the ImageJ software.

### Generation of ZAS overexpression transgenic lines in rice

The full-length cDNA of *ZAS* (*LOC_Os09g15240*) was PCR-amplified using primers *ZAS-OX-F* and *ZAS-OX-R* (table S1) and cloned into the corresponding site downstream of the *35S* promoter in the binary vector *pCAMBIA1300*, yielding the plasmid *pCAMBIA1300-ZAS*. The plasmid was introduced into Nipponbare wild type *Japonica* rice cultivar by *Agrobacterium tumefaciens* (strain EHA105)-mediated transformation as previously described (*53*). The transgenic seedlings were selected on half Murashige and Skoog (MS) medium containing 50 mg L^−1^ hygromycin and grown in GH till *T1* generation. Independent transgenic lines were selected, and twelve seedlings per transgenic lines were grown to identify homozygote lines at the *T2* generation. 40 seeds from each transgenic lines were germinated again on half Murashige and Skoog (MS) medium containing 50 mg L^−1^ hygromycin and transgenic lines that showed 90-100 % germination rate considered as homozygote lines. Then, *T2* homozygote transgenic *ZAS* overexpression plants were propagated to obtain *T3* generation seeds.

### Southern-blot analysis

Rice genomic DNA was isolated as previously described (*54*). Southern blotting was performed with the digoxigenin (DIG)-labelled PCR-amplified gene fragment of *HPT* probe as described in the DIG System Manual (Roche, Inc., Basel, Switzerland). Genomic DNA (20 μg) was digested overnight with *HindIII/BamHI* (only one cut site in the T-DNA region), fractionated on a 0.8% agarose gel by electrophoresis, and transferred onto an Amersham Hybond N^+^ nylon membrane (GE Healthcare). The 840 bp *HPT* probe was synthesized by PCR DIG Probe Synthesis Kit (Roche) using primers *HPT-DIG-F* and *HPT-DIG-R* (table S1). The blotted membrane was pre-hybridized in DIG Easy Hybridization solution (Roche) at 42°C for 2 h. Afterwards, a denatured DIG-labelled *HPT* probe was added, and hybridization was performed overnight at 42°C. After hybridization, the membrane was washed at 65°C, two times with 2xSSC and 0.1% SDS for 15 min, and two times with 0.1xSSC and 0.1% SDS for 15 min. Further, probe hybridization signal was examined by digoxigenin chemiluminescence detection.

### Photosynthesis parameters measurement

Leaf gas exchange was measured on the youngest fully expanded leaf of four-week-old rice plants using the LICOR-6800 Gas Exchange system and the 6800-12A chamber. Gas exchange was determined on a 1 x 3 cm leaf aperture or if the leaf was thinner, leaf width was measured, and gas exchange measurements were extrapolated for a 1x3 cm aperture. The following instrument settings were programmed: leaf VPD 1.5 kPa, Txchg 28°C, CO_2_ 410 ppm, flow 500 µmol s^−1^, ΔPcham 0.2 KPa, fan speed of 7000 rpm, PARi 400 µmol m^−2^ s^−1^ using the LED light source. The relative chlorophyll content was measured using SPAD-502Pus (Konica Minolta).

### Gene expression analysis by quantitative real-time PCR

Total RNA was extracted from rice tissues, using TRI-Reagent with the Direct-zol RNA MiniPrep Kit according to the manufacturer’s instruction (Zymo Research). RNA concentration, quality and integrity were checked using a NanoDrop 2000 UV-Vis Spectrophotometer (Thermo Scientific). Reverse transcription reaction was performed with the iScript cDNA Synthesis Kit, using 1 µg of total extracted RNA and following the manufacturer’s instructions (BIO-RAD Laboratories). Primers used for quantitative real-time PCR (qRT-PCR) analysis are listed in table S1. qRT-PCR was performed in a StepOne™ Real-Time PCR Systems (Applied Biosystems), using SYBR Green Supermix to monitor double-stranded DNA (dsDNA) synthesis following the manufacturer’s instructions. Relative expression levels of genes were determined using a comparative C_t_method as previously described (*55*) and the rice *Ubiquitin* (*UBQ*) gene was used as the internal control to normalize target gene expression.

### Sugar quantification

Sugars were extracted based on the protocol described by (*56*) with a slight modification. Briefly, 15 mg of freeze-dried powder in 2 mL tube (Eppendorf) was extracted by adding methanol (400 μL)/water (400 μL), and the samples were kept on the continuous shaker for one hour at room temperature. Next, 800 μL of chloroform added to the tubes and vortexed, then proceeded with 5 min centrifugation at 14000 RPM (Revolutions Per Minute). 20 μL of aliquot from the upper phase of extract was transferred to new HPLC tube and 10 μL of ribitol (10 mg/mL) added as internal standard, and the samples were evaporated to dryness under vacuum. Derivatization was initiated with 30 μL of methoxyamine hydrochloride (Sigma-Aldrich) at a concentration of 20 mg/mL in pyridine. The samples were incubated at 40°C for 1 h, and then 70 μL of *N*-methyl-*N*-trimethylsilyltrifluoroacetamide (Sigma-Aldrich) was added to the samples and kept again at 40°C for 45 minutes. The samples ran on GC–MS (Agilent HP6890) with a 5973MSD and a 10 : 1 split injector. For retention index external calibration, a mixture of alkanes, ranging from 10 to 30 carbons, was ran together with samples.

### RNAseq experiment and data analysis

Total RNA was extracted from rice roots and shoots of the *OX6* line, using TRI-Reagent with Direct-zol RNA MiniPrep Kit according to the manufacturer’s instruction (Zymo Research). RNA quality and quantity were checked using RNA Nano 6000 Assay Kit of the Bioanalyzer 2100 system (Agilent Technologies, CA, USA). Library construction, sequencing and data analysis were performed by Novogene Technology. The Illumina Novaseq platform was used for sequencing and 150 bp paired-end reads were generated. Paired-end clean reads were mapped to the Nipponbare reference genome (http://rice.uga.edu/pub/data/Eukaryotic_Projects/o_sativa/annotation_dbs/pseudomolecules/version_7.0/) by Hisat2 v2.0.5. Gene quantity was calculated by featureCounts v1.5.0-p3, and then FPKM of each gene was calculated based on the length of the gene and reads count mapped to this gene. Differentially expressed genes were identified using the DESeq2 R package (1.20.0) with the following conditions; an adjusted *P*-value ≤ 0.05 and fold change (FC) ≥ 0.3 (figs. S9, A and B). Principal Component Analysis (PCA) and heatmap of DEGs performed using Heatmaper (http://www.heatmapper.ca/expression/) to show the biological replicas consistency (figs. S9, C and D). Gene Ontology (GO) enrichment analysis of DEGs were implemented by the clusterProfiler R package via NovoMagic online platform (https://ap-magic.novogene.com/). GO terms with corrected *P*-value ≤ 0.05 were considered significantly enriched by differential expressed genes (figs. 9, E and F).

### Inorganic Anions Analysis by Capillary Electrophoresis (CE)

Anions were extracted from freeze-dried root and leaf tissues with ultrapure water using a mortar and pestle. The extracts were filtered using 0.22 µm filters and assayed by capillary electrophoresis (Agilent 7100, Agilent Technologies, Santa Clara, CA, USA). Sample injection was at 50 mbar for 5 s with +30 kV voltage and a detection wavelength of 310/20 nm. Sulfate (SO_4_^2−^), nitrate (NO_3_^−^) and phosphate (PO_4_^3−^) were analyzed through a bare fused silica capillary column with an extended light path BF3 (i.d. = 50 µm, I = 72 cm, L = 80.5 cm). Sample injection was followed by 50 mbar pressure for 4 s with −30 kV voltage and detection at the 350/380 nm wavelength. All anions were identified using pure standards. The final anion contents in each sample were calculated as µg g^−1^ DW (dry weight). For each genotype, at least four biological replicated were considered for each organ (root and shoot).

## Acknowledgments

We thank Dr. Alisdair Fernie and his lab for the hormone analysis. We thank Akmaral Serikbayeva, Abrar Felemban and Justine Braguy for helping with tissue culture. We thank Andrea Zuccolo, Alice Fornasiero and Yagiz Alagoz for their support with the whole genome sequencing analysis. We thank Vijayalakshmi Ponnakanti and Saad Hammad for technical support.

## Funding

This work was supported by baseline funding and Competitive Research Grant (CRG2020) given to S.A.-B. from King Abdullah University of Science and Technology (KAUST).

## Author contributions

S.A.B. and A.A. conceived and designed the research. S.A.B. supervised the study. I.H. generated the transgenic lines. A.A., M.J. and L.B. conducted GH and hydroponic experiments and analyzed phenotypic characterization. M.J. and A.A. performed Striga germination and rescue assays. A.A. prepared material for RNA-Seq and analyzed data. J.Y.W., K.X.L. and A.A. performed the metabolite analysis. V.M. and A.A. measured photosynthetic activities. I.B., A.A. and G.-T.C. prepared material and performed cellular level analysis. M.M. and G.V and analyzed the nutrients. V.F., T.M and L.L. analyzed mycorrhization level. N.-C.D., M.-H.L. and Y.-I.C.H. contributed to the analysis of agronomic traits. A.A generated the figures and wrote the manuscript with input from all authors. S.A.B. edited and approved the manuscript.

## Competing interests

All other authors declare they have no competing interests.

## Data and materials availability

The materials utilized in this study are accessible under an MTA agreement with the King Abdullah University of Science and Technology (KAUST). All data are available in the main figures or supplementary materials. The RNA sequencing data produced in this study shall be duly submitted to the NCBI BioProject database, which can be accessed at the following web address: https://www.ncbi.nlm.nih.gov/bioproject/

## References

1. A. R. Moise, S. Al-Babili, E. T. Wurtzel, Mechanistic aspects of carotenoid biosynthesis. Chemical Reviews 114, 164–193 (2014).

2. F. Bouvier, O. Dogbo, B. Camara, Biosynthesis of the food and cosmetic plant pigment bixin (annatto). Science 300, 2089–2091 (2003).

3. N. Nisar, L. Li, S. Lu, N. C. Khin, B. J. Pogson, Carotenoid metabolism in plants. Molecular Plant 8, 68–82 (2015).

4. H. Hashimoto, C. Uragami, R. J. Cogdell, Carotenoids and photosynthesis. Carotenoids in Nature: Biosynthesis, Regulation and Function, 111–139 (2016).

5. X. Zheng, G. Giuliano, S. Al-Babili, Carotenoid biofortification in crop plants: citius, altius, fortius. Biochimica et Biophysica Acta (BBA)-Molecular and Cell Biology of Lipids 1865, 158664 (2020).

6. J. C. Moreno, J. Mi, Y. Alagoz, S. Al‐Babili, Plant apocarotenoids: from retrograde signaling to interspecific communication. The Plant Journal 105, 351–375 (2021).

7. X. Zheng, Y. Yang, S. Al-Babili, Exploring the diversity and regulation of apocarotenoid metabolic pathways in plants. Front. Plant Sci. 12, 787049 (2021).

8. S. R. Cutler, P. L. Rodriguez, R. R. Finkelstein, S. R. Abrams, Abscisic acid: emergence of a core signaling network. Annual Review of Plant Biology 61, 651–679 (2010).

9. S. Al-Babili, H. J. Bouwmeester, Strigolactones, a novel carotenoid-derived plant hormone. Annual Review of Plant Biology 66, 161–186 (2015).

10. M. E. Auldridge, D. R. McCarty, H. J. Klee, Plant carotenoid cleavage oxygenases and their apocarotenoid products. Current Opinion in Plant Biology 9, 315–321 (2006).

11. O. Ahrazem, L. Gómez-Gómez, M. J. Rodrigo, J. Avalos, M. C. Limón, Carotenoid cleavage oxygenases from microbes and photosynthetic organisms: features and functions. International Journal of Molecular Sciences 17, 1781 (2016).

12. A. Alder, M. Jamil, M. Marzorati, M. Bruno, M. Vermathen, P. Bigler, S. Ghisla, H. Bouwmeester, P. Beyer, S. Al-Babili, The path from β-carotene to carlactone, a strigolactone-like plant hormone. Science 335, 1348–1351 (2012).

13. S. H. Schwartz, B. C. Tan, D. A. Gage, J. A. Zeevaart, D. R. McCarty, Specific oxidative cleavage of carotenoids by VP14 of maize. Science 276, 1872–1874 (1997).

14. B. C. Tan, L. M. Joseph, W. T. Deng, L. Liu, Q. B. Li, K. Cline, D. R. McCarty, Molecular characterization of the Arabidopsis 9‐cis epoxycarotenoid dioxygenase gene family. The Plant Journal 35, 44–56 (2003).

15. K.-P. Jia, J. Mi, S. Ali, H. Ohyanagi, J. C. Moreno, A. Ablazov, A. Balakrishna, L. Berqdar, A. Fiore, G. Diretto, An alternative, zeaxanthin epoxidase-independent abscisic acid biosynthetic pathway in plants. Molecular Plant 15, 151–166 (2022).

16. J. Y. Wang, I. Haider, M. Jamil, V. Fiorilli, Y. Saito, J. Mi, L. Baz, B. A. Kountche, K.-P. Jia, X. Guo, The apocarotenoid metabolite zaxinone regulates growth and strigolactone biosynthesis in rice. Nature Communications 10, 1–9 (2019).

17. J. Y. Wang, M. Jamil, P.-Y. Lin, T. Ota, V. Fiorilli, M. Novero, R. A. Zarban, B. A. Kountche, I. Takahashi, C. Martínez, Efficient mimics for elucidating zaxinone biology and promoting agricultural applications. Molecular Plant 13, 1654–1661 (2020).

18. J. Y. Wang, M. Jamil, M. G. Hossain, G.-T. E. Chen, L. Berqdar, T. Ota, I. Blilou, T. Asami, S. J. Al-Solimani, M. A. A. Mousa, Evaluation of the biostimulant activity of zaxinone mimics (MiZax) in crop plants. Front. Plant Sci. 13, (2022).

19. K. M. A. Perez, Y. Alagoz, B. Maatouk, J. Y. Wang, L. Berqdar, S. Qutub, M. Jamil, S. AlNasser, N. BinSaleh, P. Lin, Biomimetic mineralization for smart biostimulant delivery and crop micronutrients fortification. Nano Letters 23, 4732–4740 (2023).

20. V. Fiorilli, J. Y. Wang, P. Bonfante, L. Lanfranco, S. Al-Babili, Apocarotenoids: old and new mediators of the arbuscular mycorrhizal symbiosis. Front. Plant Sci. 10, 1186 (2019).

21. C. Gutjahr, M. Parniske, Cell and developmental biology of arbuscular mycorrhiza symbiosis. Annual Review of Cell and Developmental Biology 29, 593–617 (2013).

22. C. Votta, V. Fiorilli, I. Haider, J. Y. Wang, R. Balestrini, I. Petřík, D. Tarkowská, O. Novák, A. Serikbayeva, P. Bonfante, Zaxinone synthase controls arbuscular mycorrhizal colonization level in rice. The Plant Journal 111, 1688–1700 (2022).

23. A. Ablazov, C. Votta, V. Fiorilli, J. Y. Wang, F. Aljedaani, M. Jamil, A. Balakrishna, R. Balestrini, K. X. Liew, C. Rajan, ZAXINONE SYNTHASE 2 regulates growth and arbuscular mycorrhizal symbiosis in rice. Plant Physiology 191, 382–399 (2023).

24. X. Li, Q. Qian, Z. Fu, Y. Wang, G. Xiong, D. Zeng, X. Wang, X. Liu, S. Teng, F. Hiroshi, Control of tillering in rice. Nature 422, 618–621 (2003).

25. M. Umehara, A. Hanada, S. Yoshida, K. Akiyama, T. Arite, N. Takeda-Kamiya, H. Magome, Y. Kamiya, K. Shirasu, K. Yoneyama, Inhibition of shoot branching by new terpenoid plant hormones. Nature 455, 195–200 (2008).

26. Y. Wang, J. Li, Branching in rice. Current Opinion in Plant Biology 14, 94–99 (2011).

27. F. F. Barbier, E. A. Dun, S. C. Kerr, T. G. Chabikwa, C. A. Beveridge, An update on the signals controlling shoot branching. Trends in Plant Science 24, 220–236 (2019).

28. V. Gomez-Roldan, S. Fermas, P. B. Brewer, V. Puech-Pagès, E. A. Dun, J.-P. Pillot, F. Letisse, R. Matusova, S. Danoun, J.-C. Portais, Strigolactone inhibition of shoot branching. Nature 455, 189–194 (2008).

29. M. T. Waters, C. Gutjahr, T. Bennett, D. C. Nelson, Strigolactone signaling and evolution. Annual Review of Plant Biology 68, 291–322 (2017).

30. M. Jamil, B. A. Kountche, I. Haider, X. J. Guo, V. O. Ntui, K. P. Jia, S. Ali, U. S. Hameed, H. Nakamura, Y. Lyu, K. Jiang, K. Hirabayashi, M. Tanokura, S. T. Arold, T. Asami, S. Al-Babili, Methyl phenlactonoates are efficient strigolactone analogs with simple structure. J. Exp. Bot. 69, 2319–2331 (2018).

31. J. Y. Wang, S. Alseekh, T. Xiao, A. Ablazov, L. Perez de Souza, V. Fiorilli, M. Anggarani, P.-Y. Lin, C. Votta, M. Novero, Multi-omics approaches explain the growthpromoting effect of the apocarotenoid growth regulator zaxinone in rice. Communications biology 4, 1–11 (2021).

32. R. D. Ioio, F. S. Linhares, E. Scacchi, E. Casamitjana-Martinez, R. Heidstra, P. Costantino, S. Sabatini, Cytokinins determine Arabidopsis root-meristem size by controlling cell differentiation. Current Biology 17, 678–682 (2007).

33. R. D. Ioio, F. S. Linhares, S. Sabatini, Emerging role of cytokinin as a regulator of cellular differentiation. Current Opinion in Plant Biology 11, 23–27 (2008).

34. M. Kieffer, J. Neve, S. Kepinski, Defining auxin response contexts in plant development. Current Opinion in Plant Biology 13, 12–20 (2010).

35. A. Durbak, H. Yao, P. McSteen, Hormone signaling in plant development. Current Opinion in Plant Biology 15, 92–96 (2012).

36. I. Hwang, J. Sheen, Two-component circuitry in Arabidopsis cytokinin signal transduction. Nature 413, 383–389 (2001).

37. T. Kakimoto, Perception and signal transduction of cytokinins. Annual Review of Plant Biology 54, 605–627 (2003).

38. J. P. To, G. Haberer, F. J. Ferreira, J. Deruere, M. G. Mason, G. E. Schaller, J. M. Alonso, J. R. Ecker, J. J. Kieber, Type-A Arabidopsis response regulators are partially redundant negative regulators of cytokinin signaling. The Plant Cell 16, 658–671 (2004).

39. Y. Zhao, S. Cheng, Y. Song, Y. Huang, S. Zhou, X. Liu, D.-X. Zhou, The interaction between rice ERF3 and WOX11 promotes crown root development by regulating gene expression involved in cytokinin signaling. The Plant Cell 27, 2469–2483 (2015).

40. X. Cheng, H. Jiang, J. Zhang, Y. Qian, S. Zhu, B. Cheng, Overexpression of type-A rice response regulators, OsRR3 and OsRR5, results in lower sensitivity to cytokinins. Genet Mol Res 9, 348–359 (2010).

41. L. Margalha, C. Valerio, E. Baena-González, Plant SnRK1 kinases: structure, regulation, and function. AMP-Activated Protein Kinase, 403-438 (2016).

42. E. Baena-González, J. Hanson, Shaping plant development through the SnRK1–TOR metabolic regulators. Current Opinion in Plant Biology 35, 152–157 (2017).

43. J. Van Leene, D. Eeckhout, A. Gadeyne, C. Matthijs, C. Han, N. De Winne, G. Persiau, E. Van De Slijke, F. Persyn, T. Mertens, Mapping of the plant SnRK1 kinase signalling network reveals a key regulatory role for the class II T6P synthase-like proteins. Nature Plants 8, 1245–1261 (2022).

44. K. Yoneyama, X. Xie, H. I. Kim, T. Kisugi, T. Nomura, H. Sekimoto, T. Yokota, K. Yoneyama, How do nitrogen and phosphorus deficiencies affect strigolactone production and exudation? Planta 235, 1197–1207 (2012).

45. F. Wang, T. Han, Q. Song, W. Ye, X. Song, J. Chu, J. Li, Z. J. Chen, The rice circadian clock regulates tiller growth and panicle development through strigolactone signaling and sugar sensing. Plant Cell 32, 3124–3138 (2020).

46. K. Yoneyama, X. Xie, T. Nomura, K. Yoneyama, Do phosphate and cytokinin interact to regulate strigolactone biosynthesis or act independently? Front. Plant Sci. 11, 438 (2020).

47. D. Das, M. Paries, K. Hobecker, M. Gigl, C. Dawid, H.-M. Lam, J. Zhang, M. Chen, C. Gutjahr, PHOSPHATE STARVATION RESPONSE transcription factors enable arbuscular mycorrhiza symbiosis. Nature Communications 13, 477 (2022).

48. X. Dai, Y. Wang, W.-H. Zhang, OsWRKY74, a WRKY transcription factor, modulates tolerance to phosphate starvation in rice. J. Exp. Bot. 67, 947–960 (2016).

49. S. Ito, J. Braguy, J. Y. Wang, A. Yoda, V. Fiorilli, I. Takahashi, M. Jamil, A. Felemban, S. Miyazaki, T. Mazzarella, Canonical strigolactones are not the major determinant of tillering but important rhizospheric signals in rice. Science Advances 8, eadd1278 (2022).

50. M. Jamil, J. Y. Wang, L. Berqdar, Y. Alagoz, A. Behisi, S. Al-Babili, Cytokinins as an alternative agent for the suicidal germination Striga control strategy Weed Res. Under process, (2023).

51. J. Braguy, M. Ramazanova, S. Giancola, M. Jamil, B. A. Kountche, R. A. Y. Zarban, A. Felemban, J. Y. Wang, P.-Y. Lin, I. Haider, M. Zurbriggen, B. Ghanem, S. Al-Babili, SeedQuant: A deep learning-based tool for assessing stimulant and inhibitor activity on root parasitic seeds. Plant Physiology 186, 1632–1644 (2021).

52. J. Y. Wang, G.-T. E. Chen, M. Jamil, J. Braguy, S. Sioud, K. X. Liew, A. Balakrishna, S. Al-Babili, Protocol for characterizing strigolactones released by plant roots. STAR Protocols 3, 101352 (2022).

53. Y. Hiei, T. Komari, Agrobacterium-mediated transformation of rice using immature embryos or calli induced from mature seed. Nature Protocols 3, 824–834 (2008).

54. P. Xu, B. Bao, Q. He, E. Peatman, C. He, Z. Liu, Characterization and expression analysis of bactericidal permeability-increasing protein (BPI) antimicrobial peptide gene from channel catfish Ictalurus punctatus. Developmental & Comparative Immunology 29, 865–878 (2005).

55. K. J. Livak, T. D. Schmittgen, Analysis of relative gene expression data using real-time quantitative PCR and the 2− ΔΔCT method. Methods 25, 402–408 (2001).

56. X. Zheng, J. Mi, A. Balakrishna, K. X. Liew, A. Ablazov, R. Sougrat, S. Al‐Babili, Gardenia carotenoid cleavage dioxygenase 4a is an efficient tool for biotechnological production of crocins in green and non‐green plant tissues. Plant Biotechnology Journal 20, 2202–2216 (2022).

